# A long-term ketogenic diet in young and aged rats has dissociable effects on prelimbic cortex and CA3 ensemble activity

**DOI:** 10.1101/2023.02.18.529095

**Authors:** Abbi R. Hernandez, Maya E. Barrett, Katelyn N. Lubke, Andrew P. Maurer, Sara N. Burke

## Abstract

Age-related cognitive decline has been linked to distinct patterns of cellular dysfunction in the prelimbic cortex (PL) and the CA3 subregion of the hippocampus. Because higher cognitive functions require both structures, selectively targeting a neurobiological change in one region, at the expense of the other, is not likely to restore normal behavior in older animals. One change with age that both the PL and CA3 share, however, is a reduced ability to utilize glucose, which can produce aberrant neural activity patterns. The current study used a ketogenic diet (KD) intervention, which reduces the brain’s reliance on glucose, and has been shown to improve cognition, as a metabolic treatment for restoring neural ensemble dynamics in aged rats. Expression of the immediate-early genes *Arc* and *Homer*1a were used to quantify the neural ensembles that were active in the home cage prior to behavior, during a working memory/biconditional association task, and a continuous spatial alternation task. Aged rats on the control diet had increased activity in CA3 and less ensemble overlap in PL between different task conditions than did the young animals. In the PL, the KD was associated with increased activation of neurons in the superficial cortical layers. The KD did not lead to any significant changes in CA3 activity. These observations suggest that the KD does not restore neuron activation patterns in aged animals, but rather the availability of ketone bodies in the frontal cortices may permit the engagement of compensatory mechanisms that produce better cognitive outcomes.

**Significance Statement:** This study extends understanding of how a ketogenic diet (KD) intervention may improve cognitive function in older adults. Young and aged rats were given 3 months of a KD or a calorie-match control diet and then expression of the immediate-early genes *Arc* and *Homer*1a were measured to examine neural ensemble dynamics during cognitive testing. The KD diet was associated with increased activation of neurons in the superficial layers of the PL, but there were no changes in CA3. These observations are significant because they suggest that compensatory mechanisms for improving cognition are engaged in the presence of elevated ketone bodies. This metabolic shift away from glycolysis can meet the energetic needs of the frontal cortices when glucose utilization is compromised.

## INTRODUCTION

In old age, increased activity in hippocampal CA3 is associated with worse cognitive outcomes (Yassa et al., 2011a; Maurer et al., 2017). Conversely, increased activation in the prelimbic cortex (PL) is correlated with better cognition (Hernandez et al., 2020b). While pharmacological agents have been shown to improve either PL (Banuelos et al., 2014) or CA3 function (Koh et al., 2010; Bakker et al., 2012; Bakker et al., 2015; Robitsek et al., 2015), an effective therapeutic approach for optimizing neural activity patterns across these brain regions with divergent relationships between activation and cognitive performance has not been established.

Although the mechanisms of dysfunction are distinct between CA3 and PL, glucose utilization is disrupted in both of these structures with age (Gage et al., 1984; McNay and Gold, 2001; Gold, 2004; Pandya et al., 2016; Gardner et al., 2020). Moreover, blockade of glycolysis is sufficient to promote epileptogenesis (Samokhina et al., 2017), suggesting that impaired glucose metabolism can lead to aberrant activity patterns. While these data point to the potential of targeting energy metabolism to improve cognitive aging outcomes, enhancing insulin signaling has produced equivocal results regarding benefits to brain function (Moore et al., 2013; Barini et al., 2016; Anderson et al., 2017). In fact, age-related declines in glucose transporters within the brain (Ding et al., 2013; Hernandez et al., 2018b) suggest that targeting glycolysis will likely be insufficient to overcome neurometabolic deficiencies.

Rather than attempting to restore impaired glucose metabolism, an alternate mechanism for alleviating metabolic deficits in the brain is to shift energy metabolism from glucose to ketone body utilization. Ketosis is an alternate metabolic state that uses ketone bodies from fat to compensate when glycolysis is unavailable during fasting, intense exercise, or carbohydrate restriction. Remarkably, while glucose utilization is impaired in the brains of persons living with Alzheimer’s disease, ketone body utilization appears intact (Ogawa et al., 1996; Castellano et al., 2015). Moreover, a ketogenic diet in aged animals has been shown to reverse biochemical alterations within the PL and hippocampus that are specific to the unique disruptions of each region (Hernandez et al., 2018a; Hernandez et al., 2018b). A ketogenic diet has also been shown to improve peripheral metabolism (Hernandez et al., 2020a), and enhance behavioral performance on a task that requires frontal-hippocampal interactions (Hernandez et al., 2018a). These data suggest that a ketogenic diet may normalize neural activity across CA3 and PL circuits that becomes disrupted in advanced age.

To test the hypothesis that cognitive enhancement in aged rats on a ketogenic diet is due to a normalization of neural activity across the frontal-medial temporal lobe circuit, the current study used single cell imaging of the mRNA for the immediate-early genes (IEGs) *Arc* and *Homer*1a (*H*1a) to measure ensemble dynamics in PL and CA3 in relation to behavior. *Arc and H*1a mRNA are transcribed immediately after neuronal activity related to attentive behavior, but intranuclear *Arc* appears within 5 min, while *H*1a is detected only after 30 min, due to large introns atypical of IEGs (Vazdarjanova et al., 2002). Cytoplasmic accumulation of these IEGs is similarly staggered, with cytoplasmic *Arc* appearing 30 min after activity (Guzowski et al., 1999) whereas cytoplasmic *H*1a is not detectable until after 60 min (Bottai et al., 2002; Marrone et al., 2008). Notably, the co-expression of *Arc* and *H*1a is high, with >95% of neurons positive for cytoplasmic *Arc* also showing nuclear *H*1a. Thus, monitoring *Arc* and *H*1a transcription permits the detection of activity during three distinct episodes at approximately 60 min, 30 min and immediately prior to sacrifice (Marrone et al., 2008). In the current experiment, young and aged rats performed two different behavioral tasks for 5 min each separated by 20 min and cytoplasmic *H*1a was used to infer activity levels while the rats were resting in the home cage prior to doing behavior (baseline). This enabled the evaluation of ensemble overlap across behavioral states (active versus rest) and tasks.

## METHODS

### Subjects and Handling

Fisher 344 x Brown Norway F1 (FBN) Hybrid male and female rats from the National Institute on Aging colony at Charles River were used in this study, as these rats are a robust model of non-pathological aging (McQuail and Nicolle, 2015; Hernandez et al., 2020). 36 total young (4-7 months) and aged (20-23 months) rats were split across control (n = 6 young male, n = 8 aged male, n = 1 young female and n = 2 aged female) and ketogenic (n = 8 young male, n = 8 aged male, n = 1 young female and n = 1 aged female) diet groups. Additional females were unavailable for inclusion in this study, which precluded the analysis of any potential sex differences. The data from the female rats, however, did not vary from the means of the male rats and were thus included in the analysis. All rats were housed individually and maintained on a 12-h reversed light/dark cycle with all behavioral testing occurring in the dark phase.

### Diet

All rats were fed either a high-fat, low-carbohydrate KD (75.85% fat, 20.12% protein, 3.85% carbohydrate; Lab Supply; 5722, Fort Worth, Texas) or a calorically and micronutrient comparable control diet (CD; 16.35% fat, 18.76% protein, 64.89% carbohydrate; Lab Supply; 1810727, Fort Worth, Texas). The specific details on composition of the diets have been reported previously (Hernandez et al., 2018a; Hernandez et al., 2018b). The primary fat source in the KD was a >95% C8 medium chain triglyceride (MCT; Neobee 895, Stephan, Northfield, Illinois). Rats were weighed daily and given ∼51 kCal for males and ∼43 kCal for females at the same time each day for the first 12 weeks of the diet. Following a 12-week adjustment period to the diet and confirmation of sustained nutritional ketosis, the amount of food was restricted to reach an ∼15% reduction in body weight to motivate participation during behavioral testing. A third group of rats, utilized only as cage controls, were fed typical rodent chow ad libitum for the duration of the study. Access to water was available ad libitum for all animals.

### Behavioral Training & Testing

Rats were trained and tested within a figure-8 shaped maze as previously described (Hernandez et al., 2018a). Following habitation to the testing apparatus, rats were trained to alternate between the left and right arms of the maze. Once rats were sufficiently alternating (≥80% correct alternations on 2 consecutive days), the working memory/biconditional association task (WM/BAT) was introduced. Rats were presented with the same pair of objects in both arms. However, the ‘correct’ object was contingent upon the location within the maze. In the left arm, object A was correct, and was hiding a froot loop reward in a well beneath the object that could be obtained by moving the object. In the right arm, object B was correct.

Dislocation of an object to reveal the food well underneath was considered an object choice, and rats were not allowed to make a second choice if their first was incorrect. If rats failed to alternate correctly, they were not presented with objects nor given a food reward for that trial. All rats completed 20 trials per day for a minimum of 5 days/week. All behavioral testing took place in low light during the dark phase and a white noise generator was used to mitigate disruptions from background noise.

### Tissue collection and Arc/Homer1a catFISH labeling

On the final day of behavior, the testing session was comprised of two behavioral epochs, counterbalanced across rats. Rats completed yoked trials so that young and aged rats performed a comparable number of trials and underwent a similar duration of testing. To do so, an aged rat performed as many trials as possible within 5 minutes. The next young rat completed the same number of trials, or 5 minutes total, whichever came first. Following a 20 minute rest in their home cage, rats performed the second epoch of behavior. Rats were then immediately placed into a bell jar containing isoflurane-saturated cotton (Abbott Laboratories, Chicago, IL, USA) separated from the animal by a wire mesh shield. Animals rapidly lost the righting reflex (<30 seconds), after which they were immediately euthanized via rapid decapitation. Tissue was extracted and flash frozen in 2-methylbutane (Acros Organics, NJ, USA) chilled in a bath of dry ice with 100% ethanol (∼−70°C). Tissue was stored at −80°C until cryosectioning. Prior to cryosectioning, one hemisphere from each experimental group was blocked so that tissue from different groups would be sliced together and so each slide would contain all experimental groups to control for potential variability is staining across slides. Sectioning was performed at 20 μm on a cryostat (Microm HM550) and thaw mounted on Superfrost Plus slides (Fisher Scientific). Sliced tissue was stored at −80°C until *in situ* hybridization.

*In situ* hybridization was performed to label *Arc* and *Homer*1a mRNA for catFISH (Vazdarjanova et al., 2002; Marrone et al., 2008) using commercial transcription kits and RNA labeling mix (REF #: 11277073910, Lot #: 26591021 & REF #11426346910:, Lot # 24158720; Roche Applied Science) to generate digoxigenin-labeled and fluorescein-labeled riboprobes using a plasmid template containing cDNA. Tissue was incubated in the generated probes overnight and labeled with an anti–digoxigenin-HRP conjugate or anti-fluorescein-(Ref#: 11207733910, Lot # 43500600 & REF#11426346910; LOT#45220020; Roche Applied Science). Fluorescein (Fluorescein Direct FISH; PerkinElmer, REF# 14921915, Lot #: 2587974) and Cy3 (TSA Cyanine 3 Tyramide; PerkinElmer, REF#: 10197077, Lot #: 2629833) were used to visualize labeled cells, and nuclei were counterstained with DAPI (Thermo Scientific).

### Arc & Homer1a imaging and quantification

Fluorescence microscopy on a Keyence BZX-810 digital microscope (Keyence Corporation of America, Itasca, IL) was utilized to obtain z-stacks at 1 µm increments from 3 sections per region of interest for each rat. A total of 4 regions were imaged: deep and superficial prelimbic cortex (PL) and distal (CA3b) and proximal CA3 (CA3c), for a total of 12 images per rat (see **Figure 1A**). Because of the potential for conflating CA3a for CA2, this subregion of CA3 was not analyzed. Deep and superficial layers of PL were analyzed separately due to their distinct patterns of connectivity. Specifically, superficial PL layers are more reciprocally connected to other cortical areas, such as the perirhinal cortex (Burwell, 2000; Furtak et al., 2007; Agster and Burwell, 2009), while deep layers of PL are more connected to subcortical structures such as the nucleus reuniens (Vertes, 2002; Vertes et al., 2006; Jayachandran et al., 2019). A custom ImageJ plugin (available upon request) was utilized to segment nuclei and to classify cells as having no mRNA present, being positive for *Arc, Homer1*, or both, as well as the subcellular location of the mRNA. All experimenters analyzing cellular imaging data were blind to age, diet, and behavioral task order (WM/BAT versus alternation first). As in previous studies (e.g., Maurer et al., 2017; Hernandez et al., 2018c), only fully visible cells within the median 20% of the optical planes were considered.

**Figure 1:**
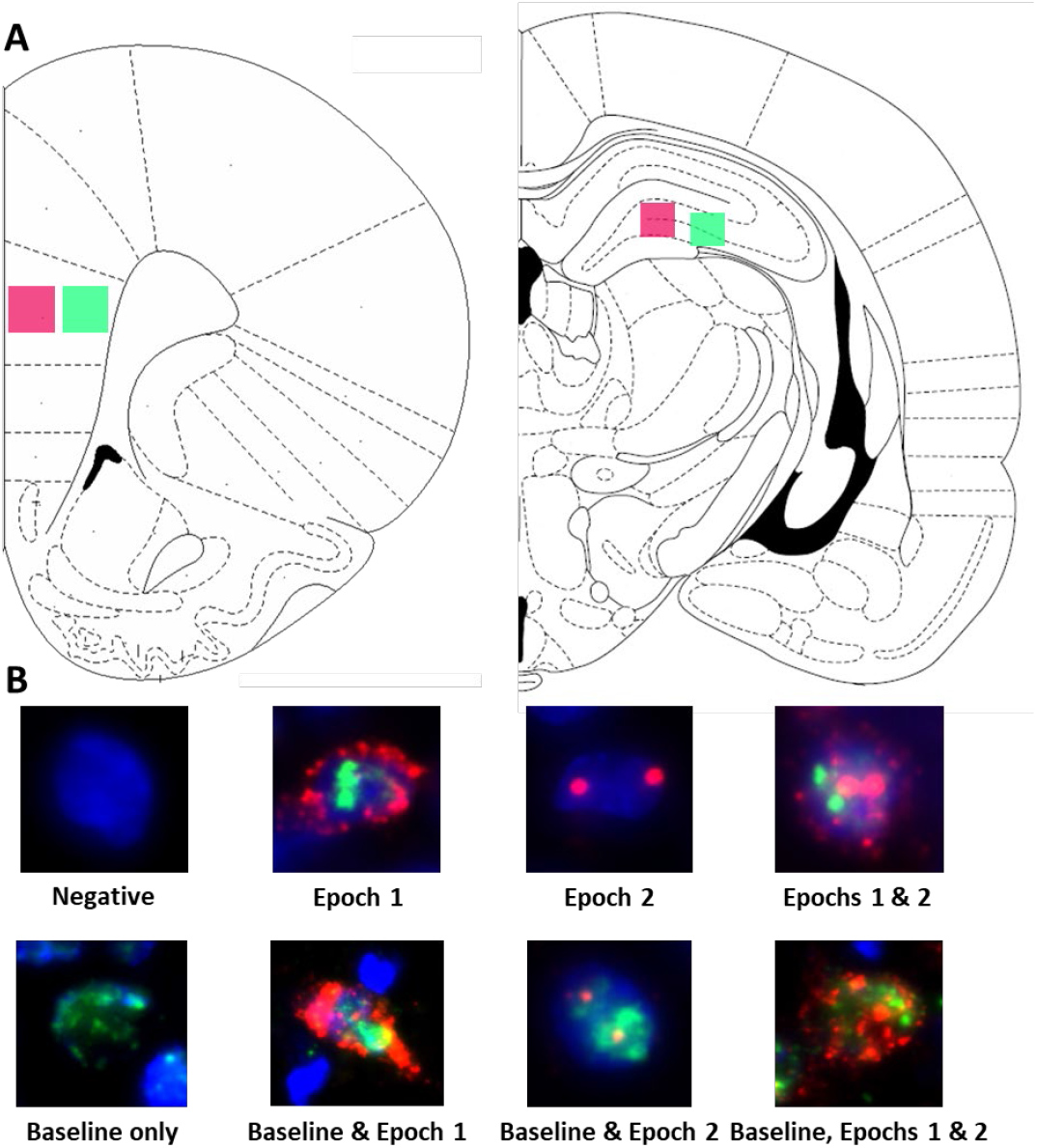
Image acquisition and analysis methodology. (A) Regions imaged within superficial (pink) and deep (green) prelimbic cortex as well as proximal (pink) and distal (green) CA3. (B) Example cells of each level of activity (see methods for further details).

**Figure 1B** shows examples of all possible cell classifications and the temporal activity profile inferred from the cellular location of *Arc* and *Homer*1a mRNA. To summarize, cells were classified as one of the following: 1) negative for any staining, indicating no activity 60 min prior to sacrifice, 2) *Arc* cytoplasm + *Homer*1a foci, indicating activity during epoch 1 only, 3) *Arc* foci, indicating activity during epoch 2 only, 4) *Arc* cytoplasm + *Homer*1a foci + *Arc* foci, indicating activity during both epochs, 5) *Homer*1a cytoplasm, indicating home cage activity only, 6) *Homer*1a cytoplasm + *Arc* cytoplasm + *Homer*1a foci, indicating home cage and epoch 1 activity, 7) *Homer*1a cytoplasm + *Arc* foci, indicating home cage and epoch 2 activity, and 8) *Homer*1a cytoplasm + *Arc* cytoplasm + *Homer1* foci + *Arc* foci, indicating activity in the home cage and during both epochs.

### Statistical analysis

Differences in neuronal activation during the behavioral epochs in relation to age, cortical layers and diet group were statistically evaluated by calculating the mean percentage of *Arc* positive cells per rat as done previously (Hartzell et al., 2013; Hernandez et al., 2018c; Hernandez et al., 2020b) to avoid inflating statistical power by including multiple measures from one animal and ensure data do not violate the assumption of independent observations (Aarts et al., 2014). Potential effects of age, layers/subregion and task were examined using repeated measures ANOVAs (ANOVA-RM) with the within-subject factors of behavioral task, cortical layers, pre-active status, and the between-subjects factors of age and diet. All analyses were performed with the Statistical Package for the Social Sciences v25 (IBM, Armonk, NY) or GraphPad Prism version 9.1.0 for Windows (GraphPad Software, San Diego, California USA, www.graphpad.com). Statistical significance was considered at p-values less than 0.05.

## RESULTS

### Home cage associated Homer1a expression in prelimbic cortex and CA3 prior to behavior

Figure 2. shows the percent of neurons that were positive for *Homer*1a in the cytoplasm, indicating activation prior to behavior during baseline for PL (**Figure 2A/B**) and CA3 (**Figure 2C/D**). The proportion of cells active at baseline was significantly different between the PL cortex and CA3 subregions (F_[3,96]_ = 44.15; p < 0.001), with more baseline activity within both subregions of CA3 compared to PL. This was not significantly affected by age (F_[1,32]_ = 1.43; p = 0.24) nor diet (F_[1,32]_ = 1.79; p = 0.19), and none of these factors significantly interacted (p > 0.24 for all comparisons).

Within the PL, there was significantly more baseline activity within the superficial layers than there was in the deep layers (F_[1,32]_ = 7.87; p = 0.008). This did not significantly vary by age (F_[1,32]_ = 0.59; p = 0.45) or diet group (F_[1,32]_ = 1.09; p = 0.31) (**Figures 2A/B**). Furthermore, there were no significant interaction effects between age, diet, and cortical layers (p > 0.18 for all comparisons). Within the deep layers, however, there were significantly fewer neurons that had *Homer*1a in the cytoplasm in aged compared to young rats (F_[1,32]_ = 5.72; p = 0.02). There was not a significant effect of age on baseline activity within the superficial layers (F_[1,32]_ = 0.03; p = 0.87).

**Figure 2:**
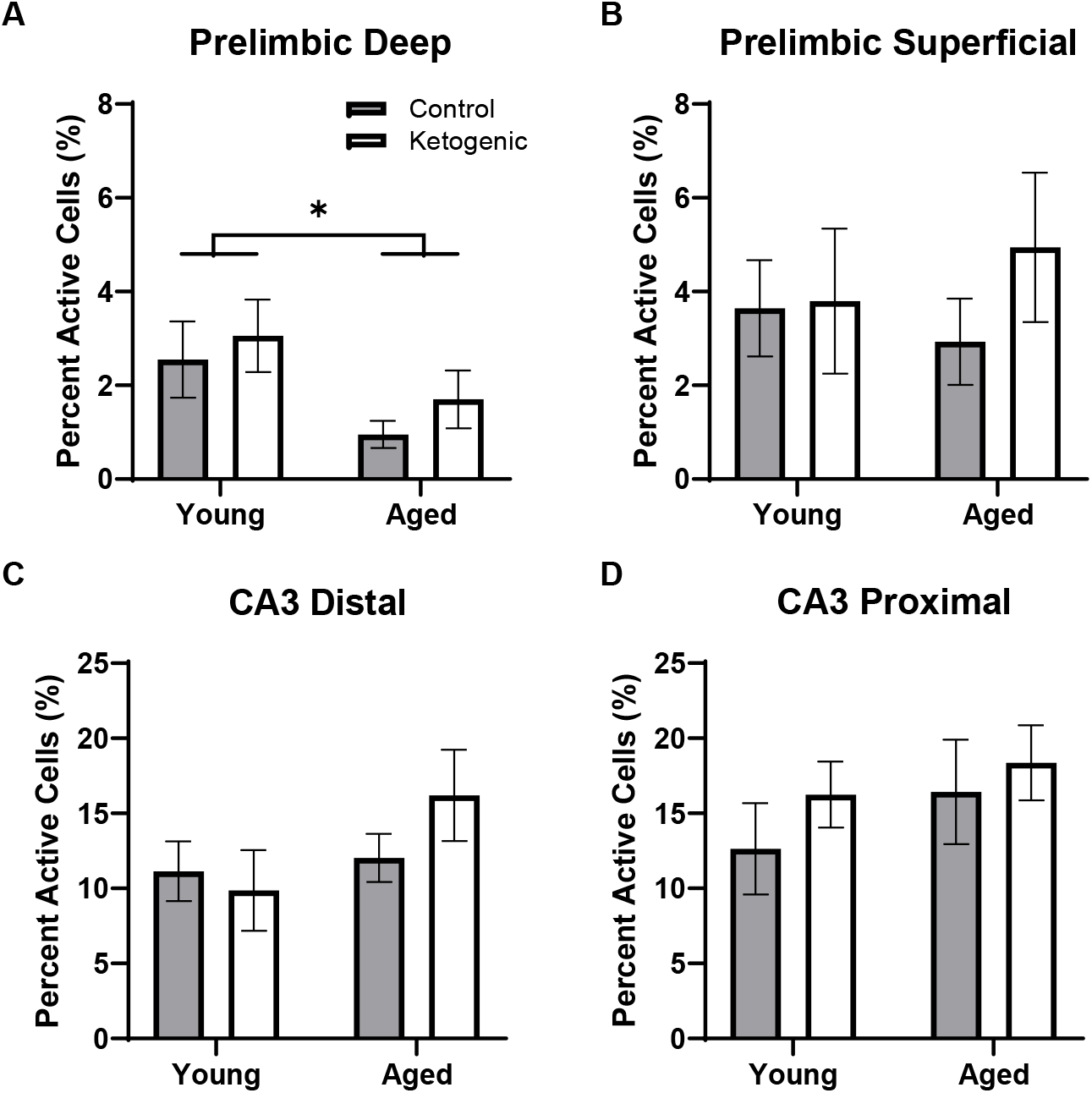
Baseline activity during home cage rest prior to behavior. (A) Aged rats had significantly decreased baseline activity within the deep layers of the prelimbic cortex. (B) There were not age or diet effects within the superficial layers of PL. (C-D) Age did not significantly influence baseline activity within distal or proximal CA3. There was no significant effect of diet on baseline *Homer*1a expression within any region or layer. Data represent group means ± 1 SEM.

The effect of subregion within CA3 on baseline activity did not reach statistical significance, though there was a trend towards greater proportion of active cells within proximal CA3 relative to distal CA3 (F_[1,32]_ = 3.68; p = 0.06; Figure BC-D). There were no differences across age (F_[1,32]_ = 2.79; p = 0.10) or diet (F_[1,32]_ = 1.15; p = 0.29) groups. Furthermore, age and diet did not significantly interact with each other (F_[1,32]_ = 0.23; p = 0.63), nor with cortical layer (p > 0.72 for both comparisons).

### Behavioral Performance and nutritional ketosis

On the final day of behavioral testing day, in which IEG expression was quantified, all rats performed two 5 min epochs of behavior separated by 20 min. In counterbalanced order, rats performed the WM/BAT task and continuous spatial alternations on the figure-8 maze. While there are age-related differences in performance accuracy on WM/BAT (Hernandez et al., 2019a) and spatial alternations (DiCola et al., 2022), all rats were previously trained to a criterion performance. Thus, on the final day of testing, there were no significant differences across age group (F_[1, 32]_ = 1.17; p = 0..29) or diet condition (F_[1, 32]_ = 0.04; p = 0.85) on WM/BAT performance accuracy, nor did these factors significantly interact (F_[1, 32]_ = 0.19; p = 0.67; **Figure 3A**). Similarly, performance accuracy for continuously alternating on the maze did not significantly differ between age (F_[1, 32]_ = 0.24; p = 0.63) or diet (F_[1, 32]_ = 0.18; p = 0.68) groups. Moreover, there was not a significant interaction between age an diet on alternation performance (F_[1, 32]_ = 0.86; p = 0.36; **Figure 3B**).

**Figure 3:**
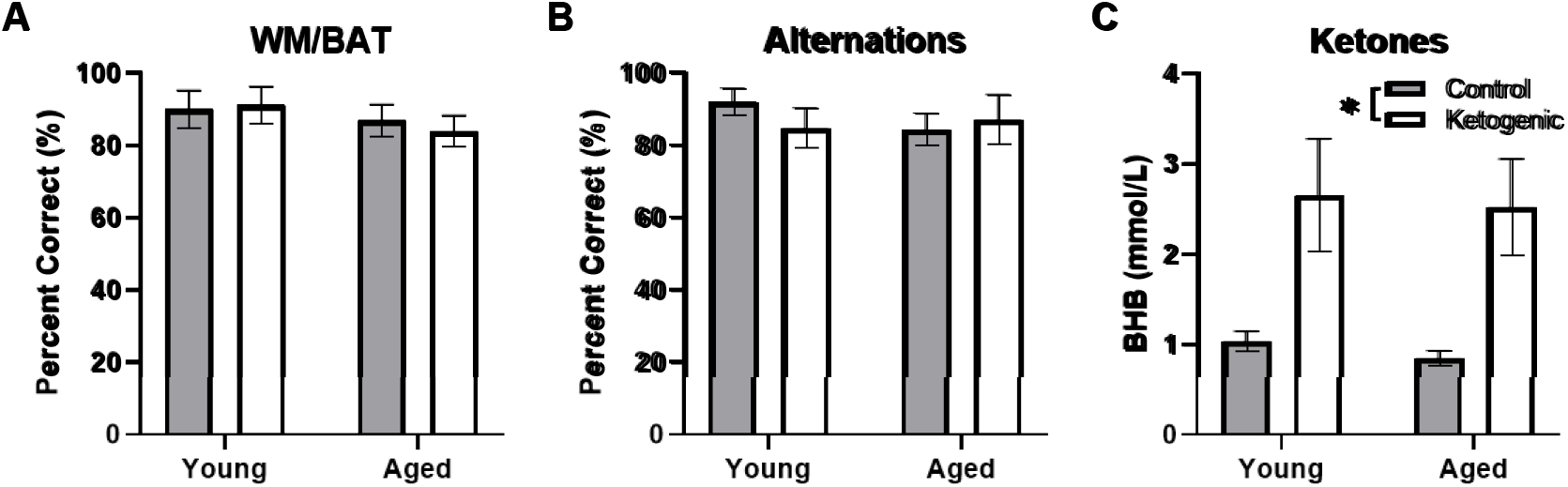
Behavioral performance and β-hydroxybutyrate (BHB) levels did not differ across age. Neither (A) WMBAT nor (B) alternation task performance were significantly affected by age or diet on the day from which cellular activity was quantified. (C) Production of β-hydroxybutyrate (BHB), a major circulating ketone body, did not differ across age groups, but was significantly elevated in rats fed a ketogenic diet relative to a control diet. Data represent group means ± 1 SEM.

Blood was collected during sacrifice to measure fasted peripheral β-hydroxybutyrate (BHB; a major circulating ketone body) immediately following behavioral testing. Rats fed a ketogenic diet had significantly more BHB than control-fed rats (F_[1,34]_ = 17.08; p = 0.0002; **Figure 3C**). There was no significant effect of age (F_[1,34]_ = 0.16; p = 0.69) nor did age significantly interact with diet (F_[1,34]_ = 005; p = 0.95), suggesting that young and aged rats had reached similar levels of ketosis.

### Age and diet influenced behaviorally-induced activity within superficial, but not deep, prelimbic cortex

**Figure 4A** shows representative images of *Arc* and *Homer*1a expression in superficial layers of PL for aged rats on a control (left panel) or ketogenic diet (right panel). The percent of cells active during the two behavioral epochs did not significantly differ between WM/BAT and the continuous alternation tasks within either the deep (F_[1,32]_ = 0.59; p = 0.45; **Figure 4B-C**) or superficial (F_[1,32]_ = 1.81; p = 0.19; **Figure 4D-E**) layers of PL. Both age (F_[1,32]_ = 13.39; p = 0.001) and the ketogenic diet (F_[1,32]_ = 17.20; p < 0.001) significantly increased activity within the superficial layers, but neither age (F_[1,32]_ = 0.52; p = 0.48) nor diet (F_[1,32]_ = 1.70; p = 0.20) significantly altered activity within the deep layers. There were no significant interactions between task, age, or diet for in with the deep or superficial layers of PL (p > 0.14 for all comparisons). The observation that a KD affected activation patterns in superficial but not deep cortical layers of PL, suggests that neurons that project to other cortical structures are more likely to be affected by interventions that modulate neurometabolism than are neurons that project to subcortical areas.

**Figure 4:**
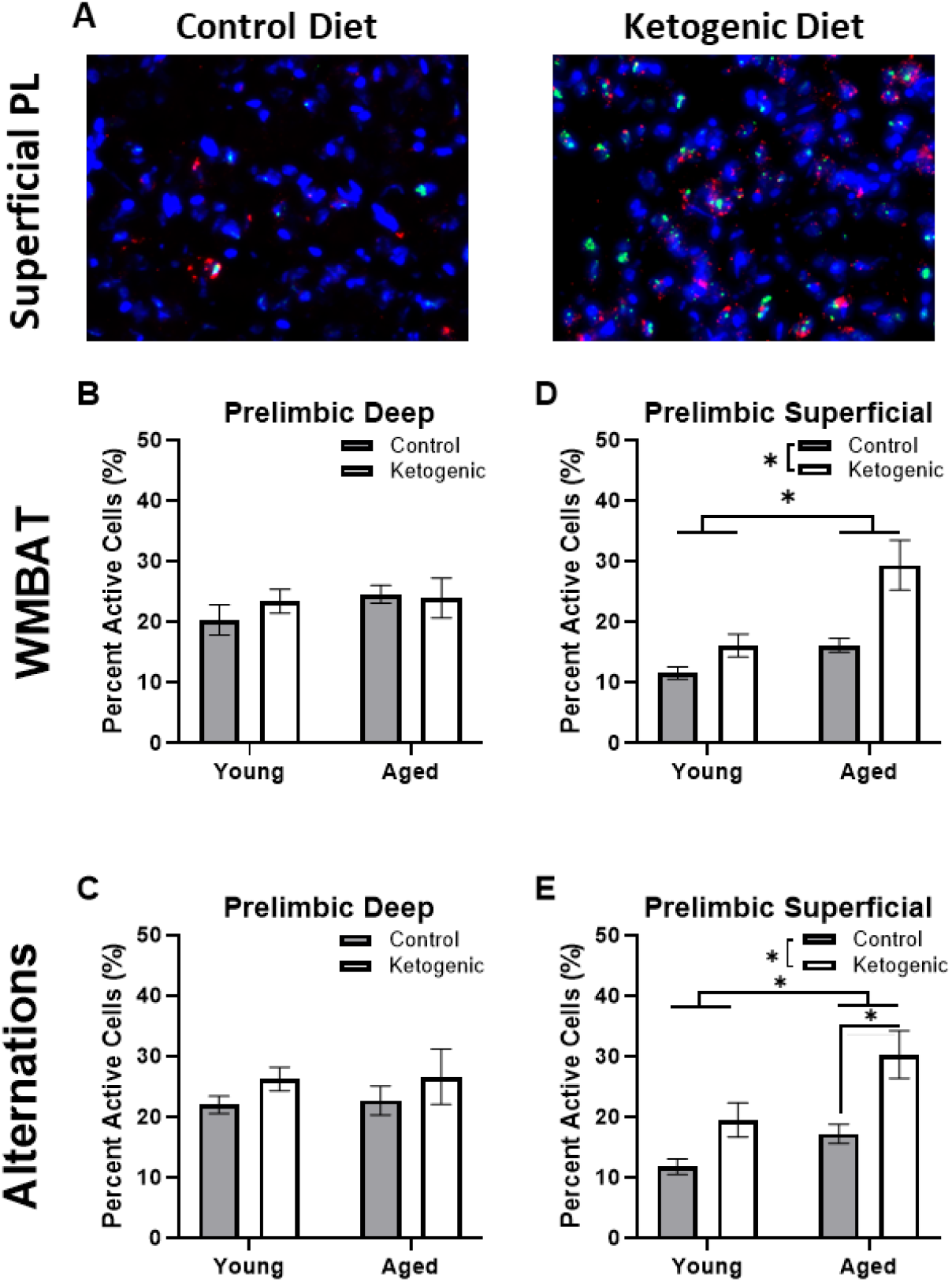
Age and diet differentially affected cellular activity within the prelimbic cortex. (A) Representative images from the superficial layer of the prelimbic cortex in aged rats on the control (left) and ketogenic (right) diets. (B-C) While there was no significant influence of age, diet nor task within the deep layers of the PL, (D-E) both age and diet significantly affected activity within the superficial layers. Data represent group means ± 1 SEM.

### Task, but not diet, influenced behaviorally-induced activity within CA3

**Figure 5A** shows representative images on *Arc* and *Homer*1a expression in distal CA3 for aged rats on a control (left panel) or ketogenic diet (right panel). In contrast to PL, the percent of cells active during the two behavioral epochs was significantly higher during WM/BAT compared to continuous alternations within both distal (F_[1,32]_ = 39.28; p < 0.001; **Figure 5B-C**) and proximal (F_[1,32]_ = 28.72; p < 0.001; **Figure 5D-E**) subregions of CA3. There was a trend for aged rats to have a higher proportion of active neurons during behavior compared to the young rats (F_[1,64]_ = 2.71, p = 0.1). However, there was no significant effect diet within distal or proximal CA3 during either task (p ≥ 0.18 for all comparisons). These observations suggest that more CA3 neurons are active when the cognitive load of the task increases.

**Figure 5:**
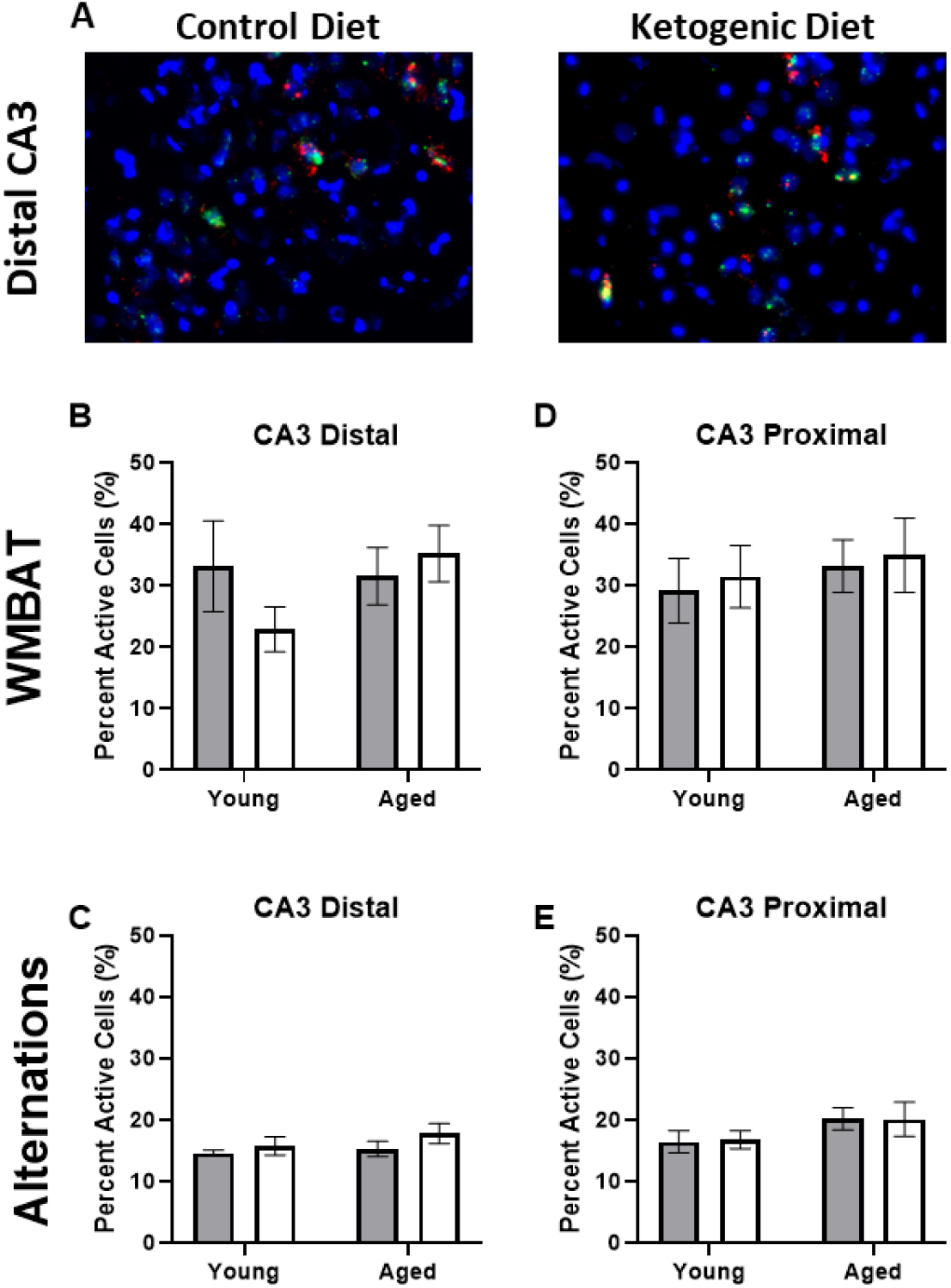
Task type influenced cellular activity within CA3. (A) Representative images from the distal portion of CA3 in aged rats on the control (left) or ketogenic (right) diets. (B-E) Activity was significantly greater during the WMBAT task relative to the alternation task within both regions of CA3 examined. However, there was no significant effect of age or diet on activity within this region. Data represent group means ± 1 SEM. (p > 0.26 for all comparisons).

### Population overlap across behavioral tasks is changed with age in prelimbic cortex but not CA3

A similarity score was calculated for each layer of the prelimbic cortex separately for each rat as done previously (e.g., Vazdarjanova and Guzowski, 2004; Burke et al., 2005; Hernandez et al., 2018c). There was no difference in similarity score across the superficial and deep layers of PL (F_[1,32]_ = 0.93; p = 0.34; **Figure 6A/B**). There was, however, a significant decrease in similarity scores in aged rats relative to the young group (F_[1,32]_ = 7.19; p = 0.01). Diet did not significantly impact similarity score (F_[1,32]_ = 0.02; p = 0.90), and there were no significant interactions effects between diet, age, and PL layers (p > 0.11 for all comparisons). These data replicate a previous observation that aged rats have reduced population overlap across two different tasks in the same spatial context compared to young animals (Hernandez et al., 2018c). Because PL neurons has been observed to fire in association with common features across different episodes (Morrissey et al., 2017), these data suggest that PL neural ensembles in older animals are less able to bridge common elements across distinct but overlapping episodes and this is not reversed by diet. In CA3, there was no difference in similarity score between distal and proximal subregions (F_[1,32]_ = 2.30; p = 0.14; **Figure 6C/D**). Furthermore, the similarity score for CA3 was not significantly affected by age (F_[1,32]_ = 0.22; p = 0.65) or diet (F_[1,32]_ = 0.46; p = 0.50). Finally, there were no significant interactions between subregion, age, or diet on similarity score

**Figure 6:**
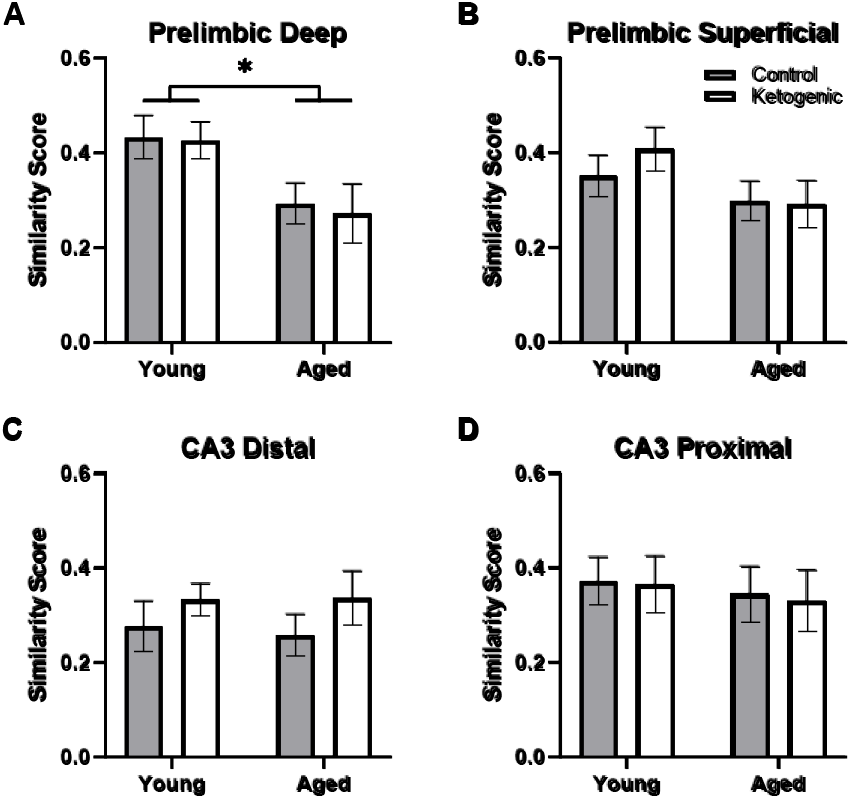
Similarity scores, representing ensemble activity overlap, were only affected by age within PL. (A) Aged rats had significantly lower ensemble activity overlap within the deep, (B) but not superficial layers of the PL. There were no differences in similarity score across diet groups in any region, nor did age significantly affect similarity in ensemble activity within (C) distal or (D) proximal CA3. Data represent group means ± 1 SEM.

### Cells with baseline activity are more likely to show activity during behavior

Previous studies have reported that neuronal populations with higher firing rates are more likely to be active across different behavioral states and contexts than cells with lower firing rates (Mizuseki and Buzsáki, 2013; Buzsáki and Mizuseki, 2014; Grosmark and Buzsaki, 2016; Witharana et al., 2016). This is hypothesized to reflect a skewed or lognormal excitability distribution of neural activity (Buzsáki and Mizuseki, 2014; Grosmark and Buzsaki, 2016). To examine the extent that age and diet may alter the skewed excitability distribution of brain organization, the behavior-related activity profiles of neurons that were active or inactive during baseline in the home cage were compared. In other words, the proportion of cells active at baseline (that is, ‘pre-active’ cells) that were also active during one or more behavioral epochs were compared to the proportion of cells inactive at baseline (that is, ‘pre-inactive’ cells) that were active during one or more behavioral epochs. Within the PL, pre-active cells were significantly more likely to have activity during behavior than pre-inactive cells (F_[1,52]_ = 123.72; p < 0.001; **Figure 7A/B**) and this behavioral-related activity bias of pre-active cells did not significantly interact with the cortical layers (F_[1,52]_ = 1.62; p = 0.21), age groups (F_[1,52]_ = 0.004; p = 0.95), or diet conditions (F_[1,52]_ = 0.20; p = 0.66). This observation suggests that both the deep and superficial layers of the PL follow a skewed excitability distribution that is not altered by age or diet. Finally, there was a significant interaction between diet and PL layer (F_[1,52]_ = 5.82; p < 0.02), such that the ketogenic diet was associated with greater activation in the superficial (F_[1,28]_ = 4.63; p < 0.05), but not deep cortical (F_[1,24]_ = 1.60; p = 0.22) layers of PL. The observation that the ketogenic diet increases activity in superficial but not deep layers of the PL is consistent with what was observed when baseline activity was not accounted for (**Figure 4**).

**Figure 7:**
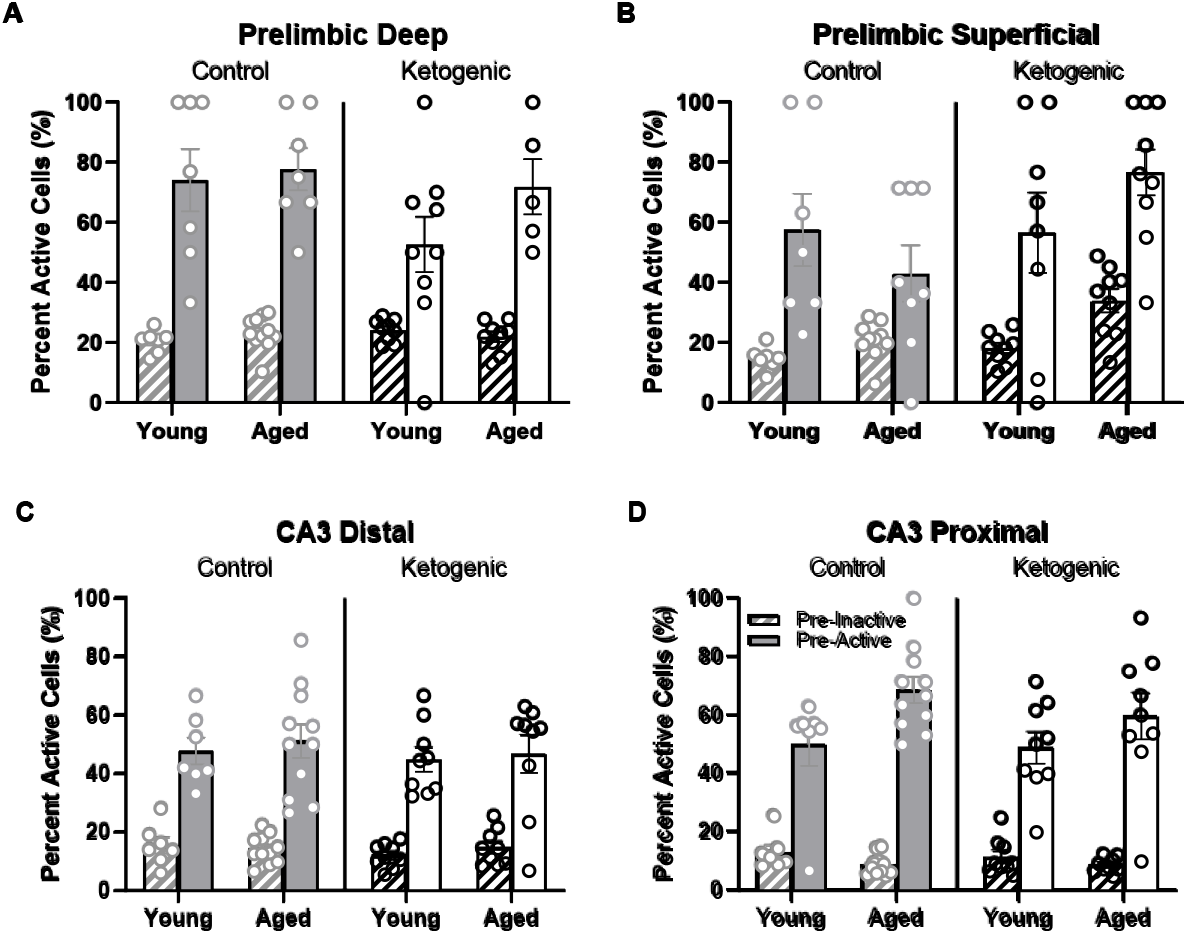
Pre-active cells were more likely to be active during behavior than pre-inactive cells. (A-B) While pre-active cells were more likely to be active within both layers of the PL, diet significantly influenced the likelihood that pre-active cells would again be active during behavior within the superficial, but not deep layers of PL. (C-D) In CA3, there was also a significant bias for pre-active to have an increased probably of also having activity during behavior compared to the pre-inactive cells. This did not vary as a function of diet condition. The pre-active bias, however, was significantly greater in proximal compared to distal CA3. Within proximal CA3, this bias was also significantly greater for young compared to aged rats.

Within distal and proximal CA3, pre-active cells were significantly more likely to have be active during behavior than pre-inactive cells (F_[1,64]_ = 360.59; p < 0.001; **Figure 7C/D**). The increased probability for pre-active cells to also show activity during behavior compared to neurons without baseline activity significantly differed between distal and proximal CA3 (F_[1,64]_ = 9.27; p < 0.003), with proximal CA3 having a larger difference in behavioral-related activity between baseline active and inactive cells than distal CA3. Different patterns of anatomical input to distal versus proximal CA3 could contribute to this observed difference in activity patterns (Witter, 2007; Lee et al., 2015; Lee et al., 2021; Lee et al., 2022). For example, distal CA3 receives more direct input from the entorhinal cortex while proximal CA3 receives mossy fiber input from both the supra- and infrapyramidal blades of the dentate gyrus (Witter, 2007). The increased probability for pre-active cells to also show activity during behavior compared to neurons without baseline activity also significantly differed between age groups (F_[1,64]_ = 5.75; p < 0.02), with older rats having a greater difference between cells that showed activity prior to behavior versus those cells that were quiescent in the home cage. In other words, aged CA3 pyramidal cells that had activity in the home cage before behavior (i.e., pre-active) were more likely to fire during behavior than were young pre-active CA3 cells. This observation is consistent with previous reports of elevated activity within aged CA3 neurons relative to young animals (Wilson et al., 2005; Thomé et al., 2016; Maurer et al., 2017; Lee et al., 2021). The behavioral-related activity bias of pre-active neurons did not significantly interact with diet (F_[1,64]_ = 0.60; p = 0.42). This suggests that aged CA3 neurons have activity dynamics that are more likely to not change across different behavioral states compared to young CA3 neurons, which is consistent with what has been observed from electrophysiological recordings (Wilson et al., 2005; Wilson et al., 2006; Lee et al., 2022). Importantly, diet condition did not have any significant effects on the pre-active versus pre-inactive bias, nor did it interact with age or CA3 subregion (p > 0.22 for all comparisons). This observation is consistent with a previous finding that the ketogenic diet did not alter the expression of genes that were related to synaptic transmission within CA3 (Hernandez et al., 2019b).

## DISCUSSION

The present study utilized the cellular compartment analysis of temporal activity by fluorescence *in situ* hybridization (catFISH) for the immediate early genes (IEGs) *Arc* and *Homer*1a (Guzowski et al., 1999; Vazdarjanova et al., 2002) to quantify neuronal activity during three distinct epochs (Marrone et al., 2008): one prior to behavior, one while rats performed the working memory/biconditional association task (WM/BAT), and another while rats performed a continuous spatial alternation task. Activity was quantified in young and aged rats fed a ketogenic (KD) or calorically equivalent control diet (CD) to test the hypothesis that a KD can reverse aberrant activity dynamics that have been associated with poor cognitive performance in the aged prelimbic cortex (PL) (Hernandez et al., 2018c; Hernandez et al., 2020b) and CA3 subregion of the hippocampus (Wilson et al., 2005; Maurer et al., 2017; Lee et al., 2021; Lee et al., 2022). Previous results were replicated regarding elevated IEG expression in aged CA3 neurons (Maurer et al., 2017), altered ensemble dynamics in PL (Hernandez et al., 2018c; Hernandez et al., 2020b), and a bias for neurons that were active in the home cage prior to behavior to also be active during behavior compared to those cells that did not show activity in the home cage (Mizuseki and Buzsáki, 2013; Grosmark and Buzsaki, 2016). The novel findings observed in the current study are that the KD increased the proportion of active neurons in the superficial layers of the PL during both behaviors in the aged rats. Within the PL, there was a reduction in ensemble overlap for neurons that active during WM/BAT and spatial alternations in aged compared to young rats, as previously observed (Hernandez et al., 2018c). This age difference was not affected by the KD, however. The KD also did not lead to any differences in activation within either distal or proximal CA3. Within CA3, the aged rats had an increase in the probability that a neuron that was active in the home cage prior to behavior would also be active during behavior compared to young rats. This was particularly evident in proximal CA3, but this age difference was not affected by the KD.

Previous studies have shown that a KD diet with medium chain triglyceride oil as the primary fat source is able to improve cognitive performance in both young and aged rats on both spatial alternations and WM/BAT (Hernandez et al., 2018a). Additionally, the KD leads to increased expression of vesicular GABA transporter in both the hippocampus and frontal cortex (Hernandez et al., 2018b), and alters the gut microbiome composition (Hernandez et al., 2022a; Hernandez et al., 2022b). Importantly, biochemical effects of the KD have been reported to differ between the hippocampus and the frontal cortex. Specifically, in the hippocampus the age-related decrease in expression of the monocarboxylate transporter (MCT) 4, which is primarily on astrocytes, is restored to the levels of young animals by the KD (Hernandez et al., 2018b). In the frontal cortex, the age-related decrease in expression of MCT2, which is primarily on neurons, is restored to the levels of young animals by the KD (Hernandez et al., 2018a). The current findings add to this growing body of evidence that a systemic KD intervention has distinct effects on the frontal cortex and the hippocampus.

CA3 undergoes a number of alterations in advanced age that have been linked to impairments on hippocampus-dependent behaviors. These include changes in synapses (Villanueva-Castillo et al., 2017; Buss et al., 2021), gene expression (Haberman et al., 2011; Hernandez et al., 2019b), altered neuron firing patterns (Wilson et al., 2005; Robitsek et al., 2015; Thomé et al., 2016; Lee et al., 2021; Lee et al., 2022), and elevated activation (Yassa et al., 2011b; Maurer et al., 2017). One previous study that used Real-Time PCR to quantify the expression of genes related to synaptic transmission and plasticity reported that 3 months of a KD diet in young and aged rats did not lead to any significant changes in gene transcription within CA3. Thus, it is conceivable that age-related changes in CA3 synaptic transmission are not reversed by a KD (Hernandez et al., 2019b). The current data are consistent with this idea and extend it to suggest that age-related increases in CA3 excitability are also not affected by a long-term KD intervention. Notably, when whole hippocampus homogenates are analyzed, 3 months of a KD does lead to increased expression of the vesicular glutamate transporter (Hernandez et al., 2018b), and decreased expression of the protein rho-associated coiled-coil containing protein kinase 2 (ROCK2) (Hernandez et al., 2019b), which regulates cytoskeletal elements to modify spine stability and is associated with synaptic loss (Huentelman et al., 2009; Swanger et al., 2015). Thus, the effects observed from whole hippocampus homogenates must be due to changes in other hippocampus subregions. The dentate gyrus in particular appears to undergo significant changes in the transcription of synapse-related genes following 3 months of a KD (Hernandez et al., 2019b). Because advanced age is associated with reduced metabolic activity (Small et al., 2002; Small et al., 2004) and lower levels of *Arc* expression in the dentate gyrus of aged compared to young animals (Small et al., 2004; Penner et al., 2011), future experiments should examine whether or not a KD can restore normal activity levels within the dentate gyrus of old animals.

While activity within CA3 was not affected by the KD, the proportion of neurons that were active during behavior increased in the KD-fed rats in the superficial layers of the PL compared to animals on the control diet. While this occurred in both age groups, the increase in superficial layer neuron activation was particularly evident in the aged rats. This observation has implications for the potential mechanisms for how a KD intervention improves cognitive performance. A previous study reported that aged rats with greater activation the of PL neurons that projected to the perirhinal cortex had better performance on WM/BAT (Hernandez et al., 2020b). Relatedly, fMRI research with human study participants has shown that higher BOLD-signal values in the frontal cortices of older adults are associated with better cognitive performance (Cabeza et al., 2002; Cabeza et al., 2004; Lighthall et al., 2014). This suggests that in the face of age-related changes in medial temporal activity, compensatory activation of frontal cortices is engaged to maintain cognitive function. Because glucose utilization in the frontal cortices is disrupted in advanced age (Gage et al., 1984; Castellano et al., 2019), but ketone body metabolism remains intact (Croteau et al., 2018; Castellano et al., 2019), it is possible that a KD increases the availability of the ketone bodies β-hydroxybutyrate and acetoacetate for neuronal metabolism. Ketone body elevation may therefore support enhanced activation in the prelimbic cortices of aged animals. In other words, dietary ketosis may increase the energy supply to the frontal cortices enabling enhanced activation that can promote compensation, leading to better cognitive outcomes. Notably, it does not appear that the KD restores age-related changes in neural ensemble activity to the patterns that are observed in young animals.

Several limitations in the current experiments are worth mentioning. First, the experiments were not sufficiently powered to consider sex as a biological variable. While a small number of females were included, the lack of availability of female rats of the Fischer344 x Brown Norway hybrid strain during the time that these experiments were conducted precluded the ability to have matched sample sizes for both sexes. Other studies have reported sex differences in neuronal metabolism with age (Yao et al., 2011; Ding et al., 2013; Yoshizawa et al., 2014; Shang et al., 2020). It is therefore critical that future research evaluate the efficacy, metabolic, and other biochemical effects of a KD in males and females. Another caveat of the current data is that both diet groups were fed only once daily and underwent modest caloric restriction to encourage motivation on the appetitive-based behavioral testing. Due to this time-restricted feeding, both groups experienced intermittent fasting. Intermittent fasting, even with a normal control diet, can affect gene expression in the brain (Castrogiovanni et al., 2018; Spezani et al., 2020; Ng et al., 2022), confer neuroprotection (Halagappa et al., 2007; Mattson et al., 2017; Anton et al., 2018), and alter the gut microbiome (Hernandez et al., 2022a; Hernandez et al., 2022b). Because all rats received intermittent fasting, it is possible that potential diet by age interactions were less than would have been observed with free feeding animals. In fact, rats the receive ad libitum feeding from middle-to old-age have worse cognitive performance compared to both ketogenic and control-fed rats that received intermittent fasting during the same time period (Hernandez et al., 2022b).

In conclusion, a KD diet was associated with increased activation of neurons in the superficial layers of the PL. This may reflect the engagement of compensatory mechanisms for improving cognition in the presence of elevated ketone bodies, which can meet the energetic needs of the frontal cortices when glucose utilization is compromised. In contrast to PL, CA3 ensemble activation patterns were not affected by diet.

## Acknowledgements

This work was supported by NIH/NICHD 2T32HD071866-06 (ARH), NIH/NIA K99AG078402 (ARH), NIH/NIA RF1AG060977 (SNB) and Florida Department of Health Ed and Ethel Moore Alzheimer’s Disease Research Program 20A016 (SNB).

## Notes

**Conflict of interest statement:** The authors declare no competing financial interests.

### Competing Interest Statement

The authors have declared no competing interest.

